# Diversity and characteristics of pTet family plasmids revealed by genomic epidemiology of *Campylobacter jejuni* from human patients in Toyama, Japan from 2015 to 2019

**DOI:** 10.1101/2022.06.28.498051

**Authors:** Daichi Morita, Hiroki Arai, Junko Isobe, Emi Maenishi, Takanori Kumagai, Fumito Maruyama, Teruo Kuroda

## Abstract

This study investigated 116 clinical isolates of *Campylobacter jejuni* from Toyama, Japan, which were isolated from 2015 to 2019. Antimicrobial susceptibility testing and whole-genome sequencing were used for phenotypic and genotypic characterization to compare antimicrobial resistance (AMR) profiles and phylogenic linkage. The multilocus sequence typing approach identified 37 sequence types (STs) and 15 clonal complexes (CCs), including 7 novel STs, and the high frequency CCs were CC21 (27.7%), CC48 (10.9%), and CC354 (9.9%). Overall, 58.6% of the isolates were resistant to at least one of the antibiotics and 3.4% were resistant to three or more antibiotic classes. The AMR profiles and related resistant factors were as follows; fluoroquinolones (51.7%), mutation in QRDRs (GyrA T86I), tetracyclines (27.6%), acquisition of *tet(O)*, ampicillin (5.2%), promoter mutation in *blaOXA193*, aminoglycosides (1.7%), acquisition of *ant(6)-Ia* and *aph(3’)-III*, chloramphenicol (0.9%), acquisition of *cat*. The resistance factors of fosfomycin (1 strain), sulfamethoxazole-trimethoprim (2 strain), and linezolid (1 strain) resistant isolates were unknown. The acquired resistance genes, *tet(O), ant(6>)-Ia, aph(3’)-III*, and *cat*, were located on pTet family plasmids. Furthermore, three pTet family plasmids formed larger plasmids that incorporated additional genes such as the Type IV secretion system.

A comparison of pTet family plasmids in Japan has not been reported, and these results imply that the diversity of pTet family plasmids has increased. The prevalence of ST4526, belonging to CC21, in Japan has been reported, and it was also the major ST type (10.9%) in this study, suggesting that the ST4526 prevalence continues in Japan.

## Introduction

*Campylobacter* is a gram-negative commensal bacterium found in the gastrointestinal tract of many animals. Particularly *Campylobacter jejuni* are among the major causes of enteritis and diarrhea in humans in many countries, including Japan (1). Based on the annual food poisoning statistics compiled by the Ministry of Health, Labour and Welfare (MHLW) in Japan, since 2003, *Campylobacter* food poisoning has become the most prevalent bacterial foodborne diseases (2). According to the Center for Disease Control and Prevention (CDC) FoodNet surveillance program in 2020 (http://www.cdc.gov/foodnet/surveillance.html), *Campylobacter* causes incidence in 14.35 per 100,000 population among the causes of laboratory-confirmed bacterial foodborne illnesses in the United States. Livestock, such as poultry, cattle and swine are asymptomatic reservoirs of pathogenic *Campylobacter*, and *Campylobacter*-contaminated food and water are major causes of foodborne illness worldwide (3, 4). In particular, consumption of poultry products is a risk factor for *Campylobacter* food poisoning (3, 4).

In addition to food poisoning, *Campylobacter* infections may cause chronic diseases such as reactive arthritis, endocarditis, cholecystitis, sepsis, and autoimmune diseases (1, 5). The most important postinfectious complication of *C. jejuni* infection is the Guillain-Barré syndrome (GBS), a demyelinating disorder resulting in acute muscular paralysis. In most studies worldwide, the mortality rate for GBS is 2-10% and *C. jejuni* infections are a common trigger of GBS (probably preceding 30% of GBS cases) (1, 6). *Campylobacter* infections are self-limiting in most of the cases and do not require antibiotics. However, antibiotic treatment using fluoroquinolones or macrolides could be used for symptoms such as high fever, bloody stools, and immunocompromised states (7). Therefore, the increase in resistant *Campylobacter* poses a threat to effective treatments.

The use of antibiotics in humans and livestock has resulted in the development of many resistant strains of bacteria, which has become a global public health problem. Drug resistant *Campylobacter* is increasing in both developing and developed countries, and especially fluoroquinolone-resistant *Campylobacter* was one of the species included on the priority list of antibiotic-resistant bacteria, created by the World Health Organization (WHO) (8). In Japan, the Japanese Veterinary Antimicrobial Resistance Monitoring System (JVARM) was established in 1999 to monitor the antimicrobial susceptibility among bacterial strains isolated from cattle, swine, and poultry on farms across Japan and has conducted surveys on resistant *Campylobacter* in livestock (9).

Whole genome sequencing (WGS) is a highly informative and discriminative approach and can comprehensively analyzes resistance factors as well as many properties such as virulence factors and phylogenetic relations. However, WGS of *C. jejuni* isolated from humans in Japan has been limited (10, 11). Although serotyping is a popular classification method for *C. jejuni*, recently, the number of untypeable strains has been increasing. Therefore, genome-based phylogenetic analysis using WGS can be used to investigate the prevalence of specific lineages. pTet family plasmids are known to be involved in resistance transmission in *C. jejuni* (12-15), but the spread of pTet family plasmids in *C. jejuni* in Japan remains unclear. The analysis of resistance transfer factors by WGS analysis is important to control the spread of resistant bacteria. In this study, 116 *C. jejuni* strains isolated from humans in Toyama, Japan between 2015 and 2019 were subjected to antimicrobial susceptibility tests and WGS to determine the prevalence of resistance and the resistance mechanism.

## Results

### Phenotypic antimicrobial resistance analysis by antibiotic susceptibility testing

Antibiotic susceptibility testing was confirmed using 24 antibiotics with the disc diffusion method (Table S1). Breakpoints were determined according to the Clinical and Laboratory Standards Institute (CLSI), and if the CLSI had no criteria, then the criteria were based on the point at which susceptible populations and clearly differentiated populations could be distinguished. *Campylobacter* exhibits intrinsic resistance to novobiocin, bacitracin, trimethoprim, rifampicin, most of *β*-lactams, vancomycin, and polymyxin/colistin (16-18). In this study, all *C. jejuni* isolates were also resistant to novobiocin, rifampicin, colistin, and some *β*-lactams, such as cloxacillin, cefmetazole and aztreonam. Except for the intrinsic resistance, fluoroquinolone (51.7%, 60/116) and tetracycline (27.6%, 32/116) resistance were frequently observed. Rarely, ampicillin (5.2%, 6/116), streptomycin (0.9%, 1/116), kanamycin (0.9%, 1/116), sulfamethoxazole-trimethoprim (1.7%, 2/116), linezolid (0.9%, 1/116), fosfomycin (0.9%, 1/116) and chloramphenicol (0.9%, 1/116) resistance were also observed (Fig. 1). No strains resistant to cefaclor, imipenem, erythromycin, streptomycin amikacin, arbekacin, and gentamicin were identified.

**Figure 1.**
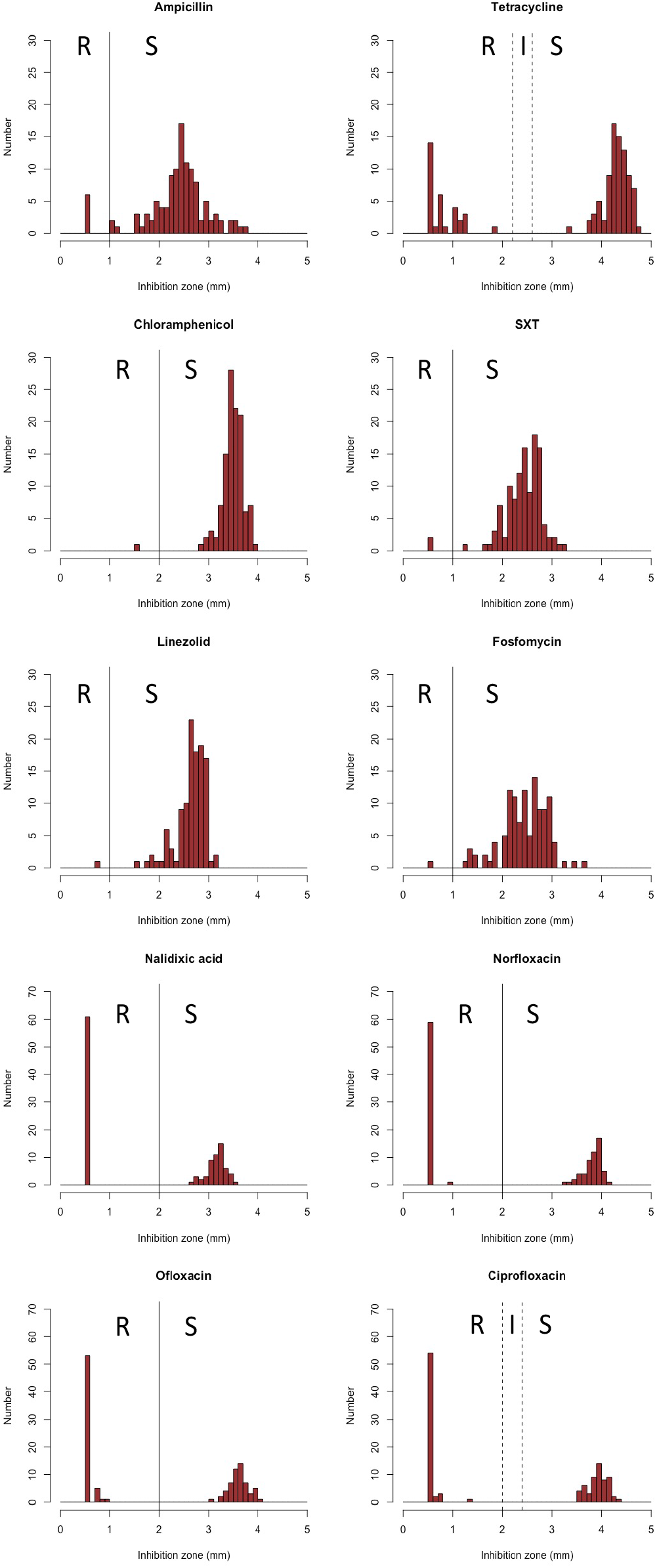
Distribution of antibiotic disc inhibition zone sizes for *C. jejuni*. Breakpoints of tetracycline and ciprofloxacin were determined according to the CLSI. In the case of antimicrobial agents for which there are no criteria in the CLSI (ampicillin, chloramphenicol, fosfomycin, linezolid, nalidixic acid, norfloxacin ofloxacin and SXT (sulfamethoxazole-trimethoprim)), the criteria were based on the point at which susceptible populations and differentiated populations distinguished.

### Genetic antimicrobial resistance analysis by whole-genome sequencing

Among 116 isolates, 101 strains were isolated within a week and subjected to WGS, except for strains with similar resistance profiles. Molecular analysis of *C. jejuni* predicted resistance factors that correlated with the phenotype corresponding to the phenotype of antimicrobial resistance (AMR) (Table S1, Fig. 2).

**Figure 2.**
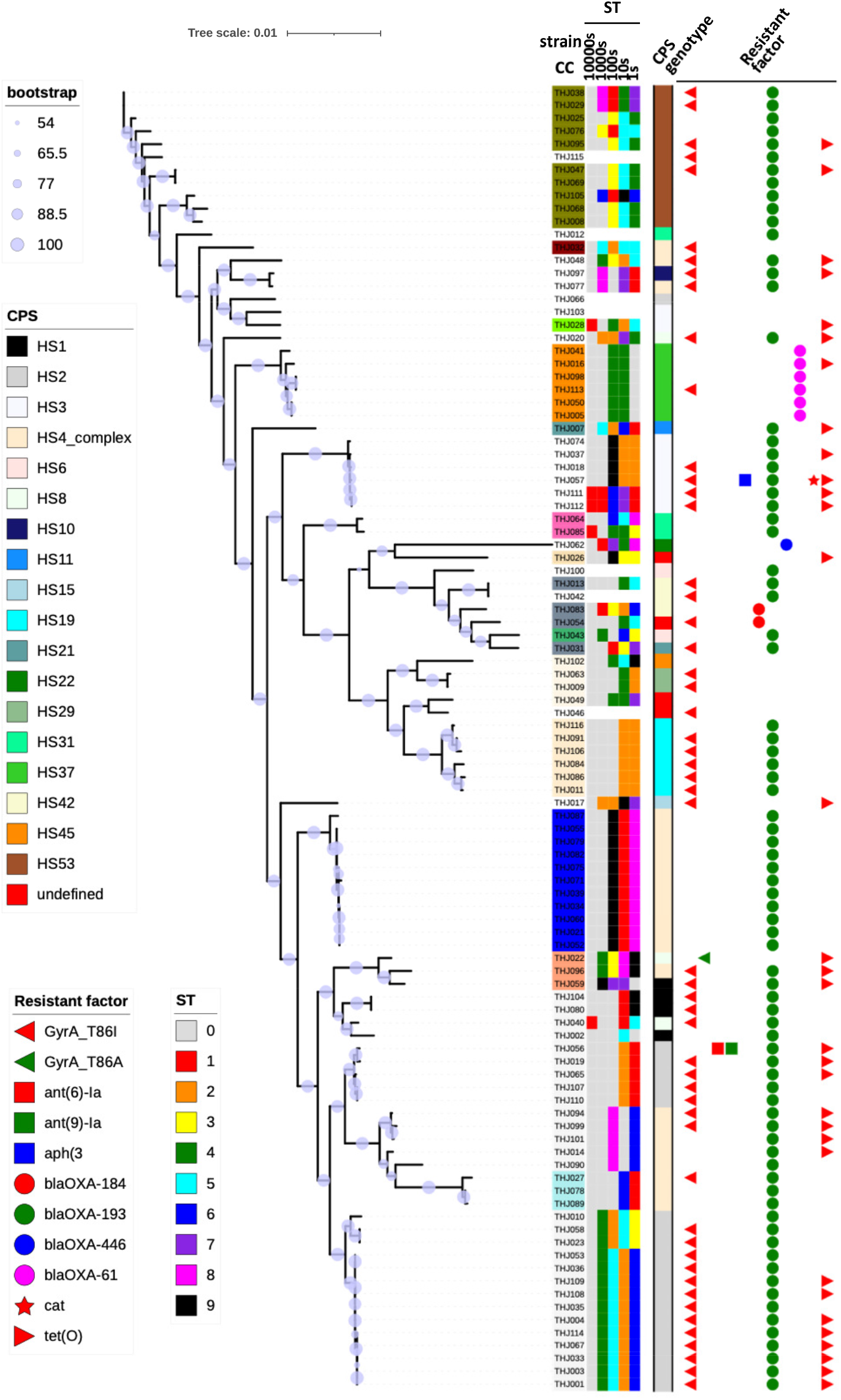
Maximum likelihood phylogenetic tree and distribution of the genotype (MLST, CPS genotype and antimicrobial resistance (AMR) genes) of the 101 *C. jejuni* isolates. Right side colored bar from inner to outer; 1st to 5th represent ST (sequence type), 6th represents CPS (capsule polysaccharide) genotyping pattern, 7th to 16th represent isolates harboring genetic determinants of resistance. The names of the isolates are color-labeled according to CC (clonal complex). The sizes of the purple circles at the nodes of phylogenetic tree indicate the magnitude of bootstrap values.

All fluoroquinolone resistant isolates were detected the mutations in quinolone resistant determining regions, (QRDRs), *gyrA* (T86I). One isolate was mutation in *gyrA* (T86A), and resistant to only nalidixic acid.

Tetracycline, streptomycin, kanamycin, and chloramphenicol resistant isolates were observed the presence of *tet(O), ant(6)-Ia, aph(3’)-III* and *cat* genes, respectively. Although the high prevalence of class D β-lactamase, *blaOXA-61, blaOXA-193*, and *blaOXA-446* were identified, only 6 ampicillin resistant isolates were found. All ampicillin resistant isolates had *blaOXA-193* and the G→T transversion in the promoter region (57 bp upstream of the annotated start codon). This mutation in the promoter has been reported to be associated with high-level expression of *blaOXA-61* in *C. jejuni* (19).

Sulfamethoxazole-trimethoprim is a combination of two antibiotics that act synergistically against a wide variety of bacteria. Although trimethoprim acts as a specific inhibitor of bacterial dihydrofolate reductase (FolA), *Campylobacter* is innately resistant to trimethoprim due to the lack of FolA (20). Therefore, this resistance may be mediated by resistance to sulfamethoxazole. In general, the sulfamethoxazole resistant mechanism has been reported as the acquisition of the low-affinity alternative dihydropteroate synthase (DHPS) genes, such as *sul1, sul2* and *sul3*, and mutations in the chromosomal DHPS gene, *folP* (21). The alternative DHPS genes *sul1, sul2* and *sul3* were not detected in the two sulfamethoxazole-trimethoprim resistant isolates, THJ056 and THJ103 Compared with the susceptible isolates, no unique mutation in *folP* up to 200 bp upstream was present in THJ103 while 27 bp insertion in *folP* was found in THJ056. It has been reported that substitution and insertion in 64^th^ Proline of FolP, such as *Escherichia coli, Neisseria meningitidis* and *Haemophilus influenzae*, are related to sulfamethoxazole resistance (22-24). The 166^th^ Proline of *C. jejuni* corresponds to the 64^th^ Proline of *E. coli*, and the amino acid insertion were located near this point, S168_R169insVYCGKEEEF (Fig. 3). In addition to the insertion, G167K mutation was also observed, and this may be responsible for sulfamethoxazole resistance.

**Figure 3.**
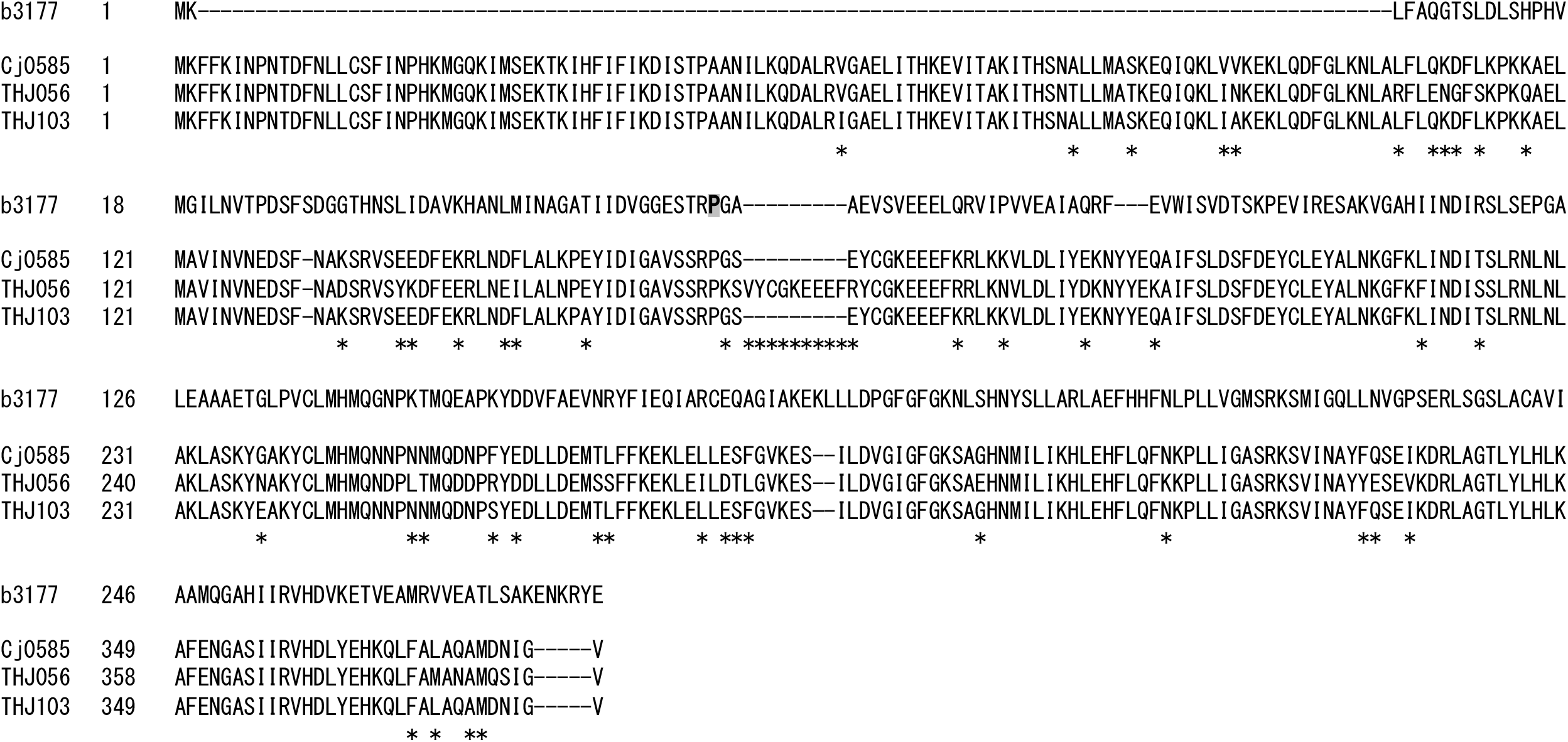
Alignment of the FolP amino acid sequences of the sulfamethoxazole-trimethoprim resistant isolates. FolP (b3177) in *E. coli* K-12 MG1655 (NC_000913.3) and FolP (Cj0585) in *C. jejuni* subsp. *jejuni* NCTC 11168 = ATCC 700819 (NC_002163.1) were used as references. FolP in sulfamethoxazole-trimethoprim resistant isolates, THJ055 and THJ105, were aligned with the references. The asterisk below the aligned sequence illustrates the amino acid substitution in FolP from *C. jejuni* NCTC 11168. Shadowed P indicates 64th Proline of FolP in *E. coli*.

In general, linezolid resistance is mediated by point mutations in domain V of the 23S rRNA (25). In addition, *optrA*, encoding an ATP-binding cassette F (ABC-F) protein that confers resistance to oxazolidinones and phenicols, has been found in *Campylobacter* (26). However, in this study, none of these resistance factors were predicted in linezolid resistant isolate.

Several mechanisms of fosfomycin resistance in gram-negative bacteria have been reported, including target modification, expression of antibiotic-degrading enzymes, reduced uptake, and rescue of the UDP-MurNAc biosynthetic pathway (27). In *Campylobacter*, the fosfomycin inactivating enzyme gene *fosX*^*CC*^ was reported (28). However, *fosX*^*CC*^ was not detected in the fosfomycin-resistant isolate.

### Molecular characterization of pTet family plasmid

The *tet(O)* gene was present in the plasmid except for one strain and the plasmid belonged to pTet plasmid family, the most prevalent plasmid type in *Campylobacter* (14, 29). Pangenome analysis showed that the plasmids had high similarity, however, a few plasmids were observed with extra DNA length that included some genes (Fig. 4). Three isolates plasmids (THJ022, THJ047 and THJ096) were similar to pCJDM67L and pCJDM202 found in *C. jejuni* and had more than 60 kbp extra DNA containing Type VI secretion system (T6SS) related genes identified in diverse species of gram-negative bacteria and acts to kill competing bacteria via a bacteriophage-like invasion and the injection mechanism (Fig. 5A) (30). The other two isolates plasmids (THJ056 and THJ057) with resistance genes other than *tet(O)* were also found similar to plasmid pGMI16-002 (CP028186.1) from the *C. jejuni* strain CFSAN054107, one was *ant(6)-Ia* and *ant(9)-Ia*, and the other was *aph(3’)-III* and *cat* (Fig. 5B) (31).

**Figure 4.**
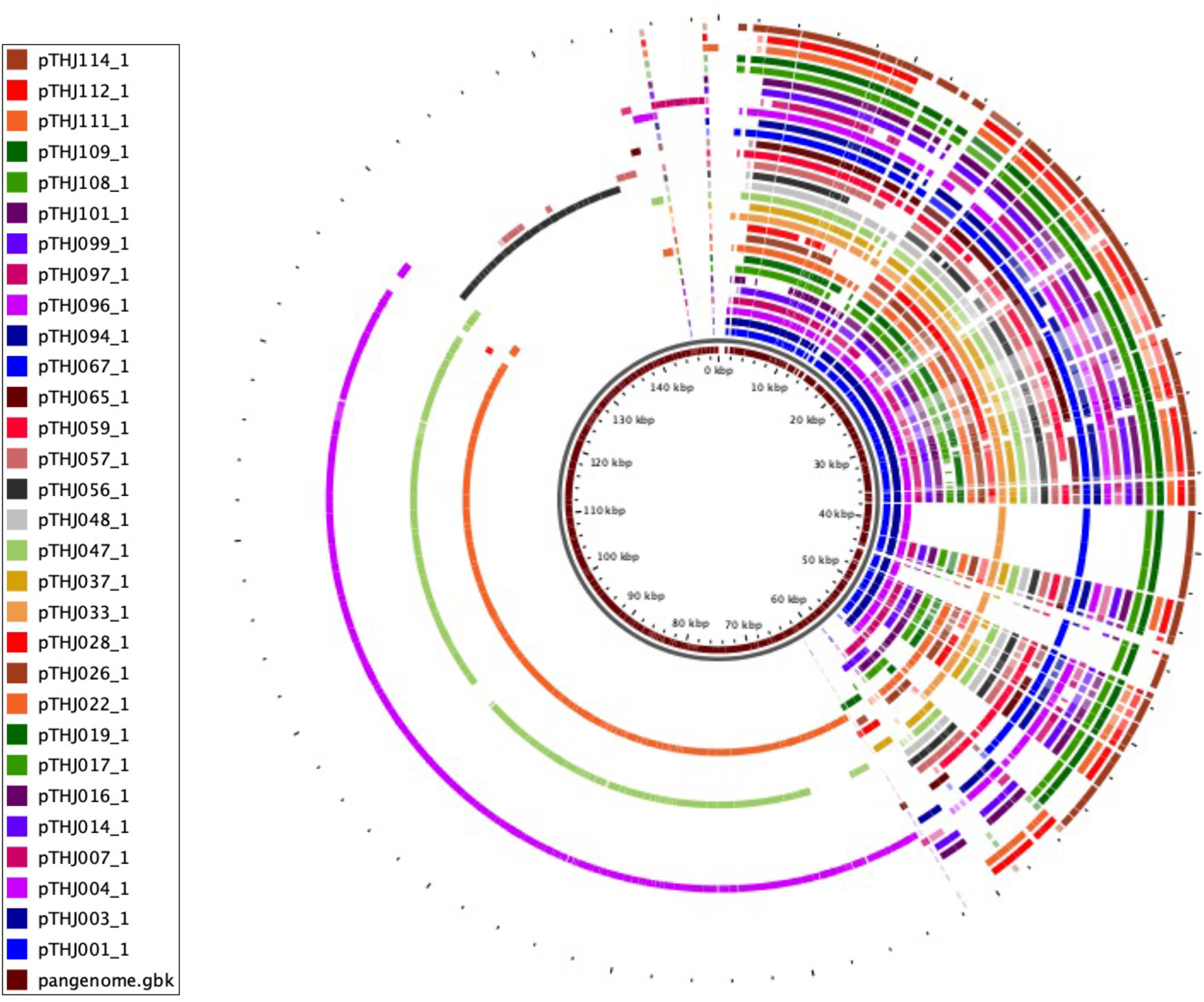
Pangenome of pTet plasmids in isolates detected by WGS. The innermost circle shows the pan-genome (brown), and outer circles indicate the plasmid.

**Figure 5.**
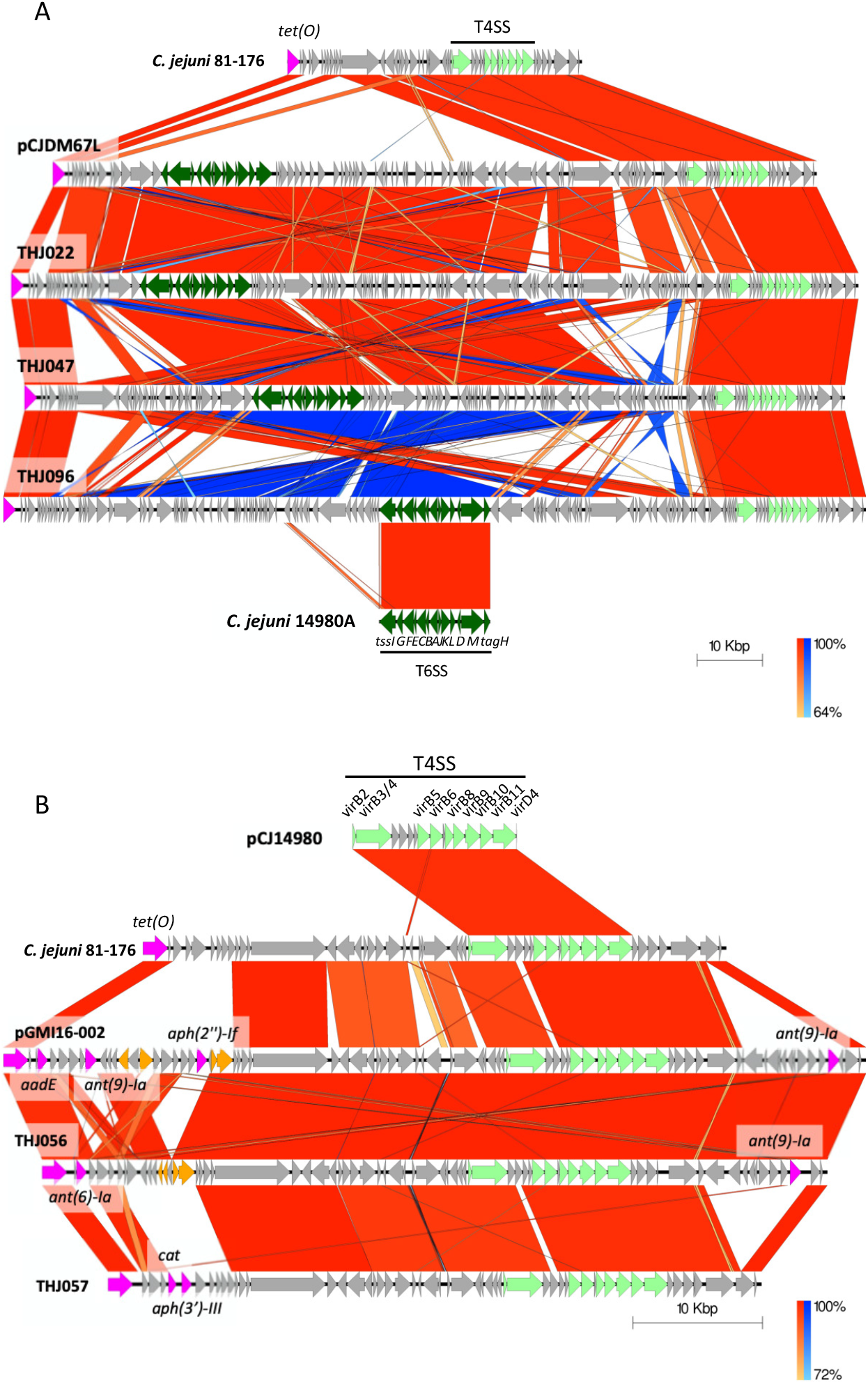
Plasmid sequence comparison. (A) Sequence alignment of pTet plasmid inserted with extra DNA containing Type VI secretion system (T6SS) related genes of isolates (THJ021, THJ047, and THJ096) with the reference pTet plasmid of *C. jejuni* 81-176 (NC_008790.1) and pCJDM67L of *C. jejuni* strain OD267 (CP014745.1). (B) Sequence alignment of pTet plasmid inserted with extra resistant genes of isolates (THJ055 and THJ056) with the reference pTet plasmid of *C. jejuni* 81-176 (NC_008790.1) and pGMI16-002 of *C. jejuni* strain CFSAN054107 (CP028186.1). Vertical blocks between sequences indicate regions of shared similarity shaded according to BLASTn (red for matches in the same direction or blue for inverted matches). Coding Sequences (CDS) are represented by colored arrows. CDSs are characterized by their functions as follows: resistance genes (pink), transposons/integrases (orange), Type IV secretion system (T4SS) genes (pale green), T6SS genes (dark green), and others (gray). The outer scale is marked in 10 kilobases. The T4SS and T6SS regions were identified by reference to *C. jejuni* strain 14980A plasmid pCJ14980 (CP017030.1) and *C. jejuni* strain 14980A chromosome (CP017029.1).

### Sequence types (STs), phylogenetic relatedness and pan-genome

Multilocus sequence typing (MLST), based on the variation among the seven housekeeping alleles defined as sequence types (STs) and clonal complexes (CCs), and is used for phylogenetic and epidemiological analyses. In the present study, the 101 isolates matched 37 STs from the reference pubMLST database, and 7 isolates did not match with the database due to an untypeable loci combination or alleles which represent novel STs. The 37 STs were linked to 15 CCs (Table S1, Fig. 2, Fig. 6).

**Figure 6.**
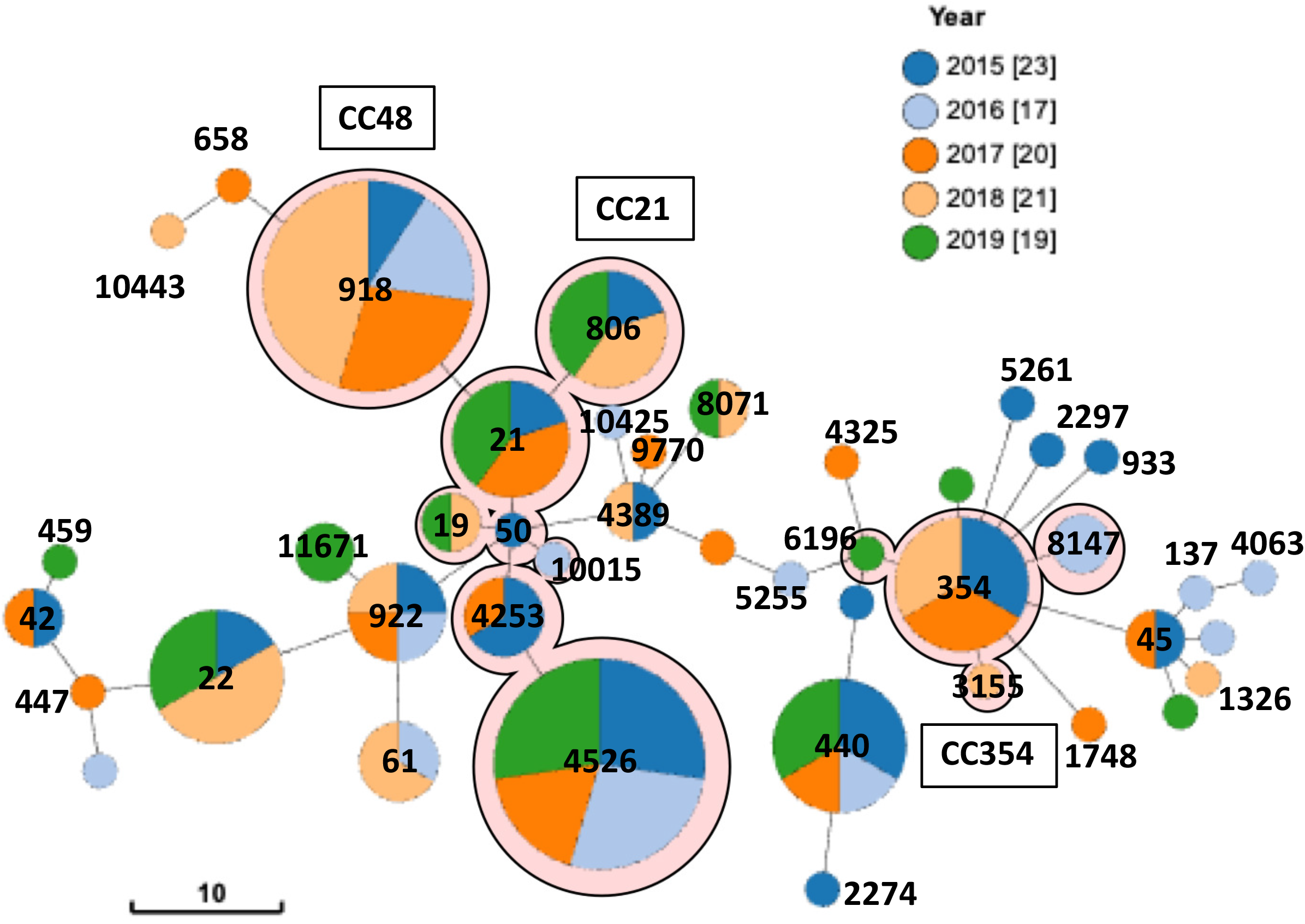
Minimum spanning tree of *C. jejuni* sequence types (STs) according to MLST profiles. Each ST is represented as a circle, with the size of the circle proportional to the number of isolates, and the inside colors indicate isolated years. The branch length represents the allelic distance. Background shading highlights typical clonal complexes (CCs) in this dataset.

The isolates were distributed widely across different CCs, with the majority belonging to CC21 (27.7 %, 28/101), CC48 (10.9 %, 11/101), and CC354 (9.9 %, 10/101). The distribution of STs throughout the years of isolation was not characterized, and there were no STs identified permanently during the five years of this study, suggesting that food poisoning by *C. jejuni* is a sporadic event in Toyama.

Comparative genomic analysis was applied for a more detailed phylogenetic analysis of the 101 isolates subjected to WGS (Fig. 2). The results of the phylogenetic analysis and MLST analysis were in good agreement. Within some STs, such as ST918, ST4526 and ST922-ST11671, genomic diversity among strains is quite low, suggesting that single clones are prevalent. The pTet family plasmid was associated with several ST types, of which 16 STs were linked to 7 CCs, and the ST types were phylogenetically distinct. Even within the same ST, not all strains possessed the plasmid, especially in ST4526, which consists of identical clones, most of the strains possessed the plasmid, but some strains lacked the plasmid. These results may indicate that plasmid acquisition occurs independently, and that acquisition and loss of plasmids occur frequently.

Penner serotyping, which is also called the heat stable (HS) serotyping has been the principal method to classify differing *Campylobacter* isolates (32). *Campylobacter* capsule polysaccharide (CPS) is the principal serodeterminant of Penner serotyping (33). Currently, 47 HS serotypes are recognized in *C. jejuni*, which correlates with the variation in CPS. Recently, HS serotyping has become untypeable in many cases, and gene-based HS classification using the PCR method for CPS loci has been proposed as an alternative method (34).

Among 116 isolates, 17 serotypes were identified, including untypeable. Although ST and HS types were correlated, half of the isolates were untypeable (n = 63) in HS serotyping, so HS serotyping was not sufficient to classify *C. jejuni* in this study (Table S1). The 101 strains that were subjected to WGS were analyzed for the regions amplified by gene-based typing of the CPR loci, as proposed by Poly et al (Table S1, Fig. 2) (34). This typing classified most of the strains (96.0%, 97/101), including those that were untypeable in HS serotyping, and the method was consistent with HS serotyping, except in 4 isolates such as THJ054 (HS38 and undefined HS in HS serotyping and gene-based), THJ056 (HS22 and HS2 in HS serotyping and gene-based), THJ113 (HS3 and HS37 in HS serotyping and gene-based typing), and THJ116 (HS3 and HS19 in HS serotyping and gene-based typing). The predominant serotype was HS4 complex (n = 21), followed by HS2 (n = 20) and HS53 (n = 11). The gene-based typing results are consistent with comparative genome analysis, and this method can be reliable in estimating the HS serotypes of *C. jejuni*.

## Discussion

WGS is a highly informative and discriminative approach, becoming a major tool in the investigation of the epidemiology in bacteria. WGS of *C. jejuni* isolated from humans in Japan has been limited and resistance factors and phylogenetic analysis were determined (10, 11). Added to these in this study, a comparative analysis of pTet family plasmids, the major resistance transfer factors of *C. jejuni*, revealed their diversity.

In this study, the major AMR in *C. jejuni*, excluding intrinsic resistance, was 51.7% for fluoroquinolones resistance and 27.6% for tetracycline resistance. Asakura et al. showed that the resistance rates of *C. jejuni* in humans and chickens were 36% for ciprofloxacin and tetracycline in 2005-2006 and were 64% for ciprofloxacin and 42% for tetracycline in 2010-2011 (35). Yamada et al. revealed that the ciprofloxacin resistance rates in the two periods 2000-2008 and 2009-2017 were 34.9% and 41.9%, respectively (36). The results of our study were in agreement with these previous studies. The main mechanisms of quinolone resistance involving *C. jejuni* have been well studied. Among them, mutations of QRDRs, in the DNA gyrase (consisting of GyrA and GyrB) and topoisomerase IV (consisting of ParC and ParE), are the targets of quinolones, is the most common and is found in almost all microorganisms. *C. jejuni* has been reported to lack the *parCE* (37-40), and a T86I in GyrA confers a high level of resistance to fluoroquinolones (41). It has also been reported that T86A in GyrA is sensitive to fluoroquinolones but confers resistance to nalidixic acid (42). In this study, all the high-level quinolone-resistant strains had T86I in GyrA, whereas the low-level quinolone-resistant strains had T86A in GyrA, which is consistent with previous studies.

The tetracycline resistance was due to the acquisition of *tet(O)*, as in previous reports of tetracycline resistant *C. jejuni*, and was mostly due to the acquisition of pTet family plasmids (12-15). The pTet family plasmids were highly homologous, but some had inserted additional sequences and expanded the resistance spectrum by the acquisition of AMEs and *cat*. Furthermore, three pTet family plasmids were megaplasmids, over 100 kbp, which incorporated additional genes such as the T6SS. Acquisition of T6SS in pTet family plasmids has been reported in several cases of *C. jejuni* and *C. coli*, and this plasmid acquisition was reported to increase *in vitro* hemolytic activity (30). Although few such plasmids were reported, the insertion of new resistance genes into pTet family plasmids, which are widely harbored in *C. jejuni*, suggests that pTet family plasmids may enhance the multidrug resistance and facilitate survival and virulence of *C. jejuni*.

*C. jejuni* is inherently resistant to most *β*-lactams but is susceptible to some *β*-lactams such as ampicillin. In this study, *blaOXA* was detected in almost all strains, but ampicillin resistance was confirmed in only a few. In the ampicillin-resistant strains, a mutation was found in the promoter region, as in previous reports, suggesting that this promoter mutation increased the expression of *blaOXA* (19). Most *C. jejuni* harbor *blaOXA*, suggesting that they are potentially resistant to ampicillin.

On the other hand, macrolide resistant strains have been reported as rare in Japan (10, 35, 36, 43), and no macrolide resistant strains were found in this study, suggesting that macrolide resistance remains rare in Japan.

The 101 isolates were classified into 37 STs, including 7 novel STs, indicating that clinical isolates of *C. jejuni* exhibit high molecular diversity. CC21 was the most common CC type (28.7%, 29/101). Among CC21 isolates, ST4526 was predominant (11/29) and showed high resistance rates to fluoroquinolones (11/11) and tetracycline (8/11). ST4526, which was first found in Japan in 2012, is a Japan-specific lineage. Oishi et al. also reported that ST4526 was the most dominant among human and chick isolates in Japan from 2007 to 2014 (11). Furthermore, Yamada et al. revealed that ST4526, while not isolated in 2000 and 2008, became the dominant ST in 2017 (36). These results suggest that ST4526 is still spreading in Japan. CC48, the second most common CC type, was detected only in ST918, which is often isolated from poultry in Japan, and was equal in number to ST4526.

In the serotyping of *C. jejuni*, the Lior method based on thermophilic antigens and the Penner method based on thermostable antigens are internationally recognized (32, 44). The major antigenic determinant in the Penner serotyping method is CPS, a cell surface molecule that affects bacteriophage infectivity (45), colonization of chickens (46), invasion of human epithelial cells (47, 48), and host immune responses (18). The CPS gene cluster contains multiple phase-variant genes interrupted by poly-G tracts, resulting in the unstable phenotype of the Penner serotype due to the on/off of expression of CPS genes due to phase variation. For this reason, Penner serotyping frequently results in untypeable strains. Recently, a method to determine the Penner genotype by PCR typing of CPS gene clusters (Penner genotyping method) was reported (34). The genotyping method was able to type most of the strains that were untypeable by the current serotyping method. In addition, phylogenetic classification by core genome analysis correlated well with the genotyping, supporting the reliability of the genotyping results. The genotyping method is considered to be an effective alternative to the Penner serotyping method because of its high typing rate and correlation with existing serotyping methods, although some issues need to be investigated further such as the expression of CPS genes.

These results suggest that the major drug resistances of *C. jejuni* clinical isolates are fluoroquinolones and tetracycline, and macrolides remain susceptible. In addition, strain ST4526 belonging to CC21 was highly resistant to fluoroquinolones and tetracycline, suggesting that it is mainly spreading in Japan. Antimicrobial susceptibility testing and molecular analysis of *Campylobacter* clinical isolates in Japan have been limited so far, and our results are consistent with previous studies. Moreover, the acquisition of resistance genes to aminoglycosides and chloramphenicol and virulence genes such as T6SS was confirmed in a part of the pTet family plasmid transmitting tetracycline resistance, and the expansion of antimicrobial-resistant *C. jejuni* is expected in the future. It is important to monitor the spread of *C. jejuni* resistance by antimicrobial susceptibility and molecular analysis. Penner serotyping, which is a typical serotyping method for *C. jejuni*, is now untypeable in many cases, and Penner genotyping is a good alternative because it can be typed in most cases, has high concordance with serotyping, and is highly correlated with phylogenetic classification by core genome analysis.

## Methods

### Bacterial isolates

*C. jejuni* strains were isolated from patients with gastrointestinal symptoms and identified by the Toyama Institute of Health in Toyama, Japan between 2015 and 2019 (Table S1). A total of 116 isolates were used, 112 of which were isolated at one hospital (JA Toyama Kouseiren Takaoka Hospital) in Toyama Prefecture, and 4 of which were collected by the Toyama City Public Health Center from a food poisoning outbreak in Toyama Prefecture. The strains were stored at -80°C in brain heart infusion broth containing 15% glycerol. Among the 112 isolates from the hospital, 26 were isolated in 2015, 19 were isolated in 2016, 22 were isolated in 2017, 28 were isolated in 2018, and 17 were isolated in 2019. Among the 4 isolates from the Public Health Center, 1 was isolated in 2016 and 3 were isolated in 2019. All strains were evaluated for antimicrobial susceptibility using the disc diffusion method, and 101 strains were subjected to WGS.

### Antimicrobial susceptibility testing

The disc diffusion methodology was based on the National Committee for Clinical Laboratory Standards (49). The disc content was as follows: ampicillin, 10 μg; cloxacillin, 1 μg; cefaclor, 30 μg; cefmetazole, 30 μg; imipenem, 10 μg; aztreonam, 30 μg; tetracycline, 30 μg; erythromycin, 15 μg; streptomycin, 10 μg; kanamycin, 30 μg; amikacin, 30 μg; arbekacin 30 μg; gentamicin, 10 μg; nalidixic acid, 30 μg; norfloxacin, 10 μg; ofloxacin, 5 μg; ciprofloxacin, 5 μg; sulfamethoxazole-trimethoprim, 23.75/1.25 μg; linezolid, 30 μg; colistin, 10 μg; chloramphenicol, 30 μg; fosfomycin, 50 μg; rifampicin, 5 μg; novobiocin, 30 μg. All the discs were Sensi-Disc (Becton Dickinson). The isolates were grown on TSA II 5% Sheep Blood Agar (Becton Dickinson) at 42°C for 24 h under microaerobic conditions (AnaeroPac-MicroAero; Mitsubishi Gas Chemical Company, Inc., Japan) Antibiotic sensitivity testing was conducted using Mueller Hinton Agar with 5% Sheep Blood (Becton Dickinson) and incubated at 42°C for 24 h under microaerobic conditions.

### Whole-genome sequencing, genome assembly, and antimicrobial resistance analysis

DNA was extracted from single-colony cultures (Monarch Genomic DNA Purification Kit, NEB). Short read libraries were prepared based on the NEBNext Ultra II FS DNA Library Prep with Sample Purification Beads (NEB) and sequenced using Illumina Miseq v3 (150 cycles). Long read libraries were prepared based on the Rapid Barcoding Kit (Nanopore) and sequenced using Oxford Nanopore MinION.

Paired-end 150 bp FASTQ files were passed through the “Bactopia 1.7.1” workflow to assess data quality, assemble contigs, and call *in silico* MLST (50). In long-read assembly, the long-read with greater than 400x coverage were randomly downsampled to a coverage of 400x using rasusa v0.6.1 (parameters: --coverage 400 --genome-size 1.7 Mb) (51). The long-reads were filtered using NanoFilt v2.8.0 (-l 3000 –headcrop 75) (52). Flye v2.9 (53) was used, and the assembled sequences were polished using Illumina reads with Pilon v1.24 (54) by default parameters. Plasmids constructed by tandem assembly of identical small plasmid sequences were manually resolved.

The assembled genomes were then passed through a bioinformatic pipeline using by BLAST techniques to identify AMR genes and AMR associated point-mutations using the programs, STARAMR 0.7.2 (https://github.com/phac-nml/staramr) and ABRicate. STARAMR databases used Pointfinder v050218 (database date on 07 Jan 2022), Resfinder v050218.1 (database date on 06 Oct 2021), MLST v2.9 (https://github.com/tseemann/mlst), and Plasmidfinder database (database date on 07 Jan 2022) (55-58). ABRicate (https://github.com/tseemann/abricate) databases used ResFinder, NCBI, CARD, and MEGARes databases (last updated on 2022-Feb-3) (58-61). FolA alignment was performed using the online version of MAFFT version 7 (https://mafft.cbrc.jp/alignment/server/) (62, 63). Minimum spanning trees were generated and visualized in GrapeTree (64).

### Plasmid comparison analysis

Pangenome analysis for plasmids from our laboratory was conducted using the GView server (https://server.gview.ca/) (65). In the GView server, blast analysis (nucleotide) was carried out using GenBank files of plasmid sequences with an e-value < 1e-10, alignment length cutoff value of 100, and percent identity cutoff value of 80. The comparison of plasmids in which gene insertion was detected with *C. jejuni* 81-176 plasmid pTet (NC008790.1) as reference was visualized using Easyfig 2.2.5 (66).

### Genome comparison and phylogenetic analysis

To identify the genus core genome, we used Panaroo v1.2.9 (67) to generate a gene presence-absence matrix with the default settings and the -a core flag to generate a core gene alignment. A core gene phylogeny was constructed from the core gene alignment using the IQ-Tree software 2.0.3 (68) with the GTR substitution model and ultrafast bootstrapping (1,000 bootstraps). Phylogenetic tree visualization was conducted using the Interactive Tree of Life v6.5 (iTOL) (https://itol.embl.de) (69).

### *In silico* identification of Penner genotype

The presence of the specific CPS sequences for a particular serotype was determined by performing a local stand-alone BLAST using a database encompassing the nucleotide sequences of CPS genotyping multiplex PCR amplification region (34). Primers and their respective PCR product sizes are listed in Table S2.

## Data availability

The genome sequences analyzed in this study are available under BioProject accession number PRJDB13586. Accession numbers and BioSample identifiers (IDs) are listed in Table S3 in the supplemental material.

## Conflicts of interest

There are no conflicts of interest to declare.

## Acknowledgements

This study was supported by a grant from the Japan Society for the Promotion of Science under Grants-in-Aid for Scientific Research (KAKENHI) (grant numbers 18K19674, 18KK0436 and 20H00562) and the Japan Agency for Medical Research and Development (grant numbers 20wm0225012h0001, 21fk0108129h0502, JP20wm0225012, JP21wm0225012 and JP22wm0225012). Illumina Miseq sequencing was conducted at the Natural Science Center for Basic Research and Development, Hiroshima University.

## Author contributions

Conceptualization: DM, FM and TKur. Formal analysis: DM, HA and FM. Funding acquisition: DM, FM and TKur. Investigation: DM, HA, JI and EM. Visualization: DM and HA. Resources: JI, EM and FM. Supervision: TKur. Writing – original draft: DM. Writing – review & editing: HA, JI, EM, TKum, FM and TKur.

